# Genome of the giant panda roundworm illuminates its host shift and parasitic adaptation

**DOI:** 10.1101/2021.05.29.446263

**Authors:** Yue Xie, Sen Wang, Shuangyang Wu, Shenghan Gao, Qingshu Meng, Chengdong Wang, Jingchao Lan, Li Luo, Xuan Zhou, Jing Xu, Xiaobin Gu, Ran He, Zijiang Yang, Xuerong Peng, Songnian Hu, Guangyou Yang

## Abstract

*Baylisascaris schroederi*, a bamboo-feeding giant panda (*Ailuropoda melanoleuca*)-specific roundworm (ascaridoid) parasite, is the causative agent of baylisascariasis, which represents a leading reason for the mortality of wild giant panda populations and therefore poses a significant threat to giant panda conservation. Here we present a 293-Mb chromosome-level genome assembly of *B. schroederi* to inform its biology, including host adaptations. Comparative genomics revealed an evolutionary trajectory accompanied by host-shift events in ascaridoid parasite lineages after host separations, suggesting their potential transmissions and fast adaptations to hosts. Genomic and anatomical lines of evidence, including expansion and positive selection of genes related to cuticle and basal metabolisms, indicates that *B. schroederi* undergoes specific adaptations to survive in the sharp-edged bamboo enriched gut of giant panda by structurally increasing its cuticle thickness and efficiently utilizing host nutrients during gut parasitism. Also, we characterized the secretome and predicted potential drug and vaccine targets for new interventions. Overall, this genome resource provides new insights into the host adaptation of *B. schroederi* to giant panda as well as the host-shift events in ascaridoid parasite lineages. These findings also add to our knowledge on the unique biology of the giant panda roundworm and aid the development of much-needed novel strategies for the control of baylisascariasis and thus the protection of giant panda populations.

## Introduction

The giant panda (*Ailuropoda melanoleuca*) is an enigmatic and endangered mammalian species endemic to Western China. Unlike other bear members in Ursidae, which are carnivores or omnivores, the giant panda almost exclusively feeds on the highly fibrous bamboo but retains a carnivoran alimentary tract [1, 2]. Consequently, the giant panda exhibits very low digestive efficiency and low metabolic rates to achieve its daily energy balance [2]. Based on this physiological situation, the tract of giant pandas likely exhibits reduced nutrient digestibility and absorption and is full of undigested and sharp-edged bamboo culms/branches [3]: the former speculation explains why giant pandas spend most of each day consuming a remarkable quantity of bamboo relative to their body size [1], and the latter illustrates potential risks to the gastrointestinal system (e.g., physical damage to the stomach and guts) and tract-inhabiting organisms, including parasitic nematodes (e.g., physical pressure and damage to worm bodies) [3, 4].

The roundworm *Baylisascaris schroederi* is the only endoparasite that is consistently found in giant pandas and has been confirmed to be one of the leading causes of death in wild giant panda populations [5–7]. In nature, *B. schroederi* infection in giant pandas follows a trophic pathway from ingestion to lifecycle completion without intermediate hosts (Figure S1). The adults inhabit the intestines of giant pandas and can cause intestinal obstruction, inflammation and even host death; in addition, the larvae might disseminate into various body tissues and induce extensive inflammation and scarring of the intestinal wall and parenchyma of the liver and lungs (also known as visceral larva migrans, VLM) [4, 5, 7]. Parasitological evidence shows that *B. schroederi* has highly evolved to adapt to its host with one body size comparable to that of other ascaridoid parasites, including *A. suum* in pigs and *P. univalens* in horses [8]. Such physical adaptations to the nutrient-limited and fiber-enriched intestinal environment of giant pandas are likely related to nutritional metabolic changes and exoskeletal cuticle resistance. However, the detailed molecular mechanisms underlying these adaptation processes remain unknown. To address these concerns and strengthen efforts to control this parasite in giant pandas, we generated the 293-Mb chromosome-level genome assembly of *B. schroederi* and compared it with those of other ascaridoid species. The analysis identified a total of 16,072 nonredundant protein-coding genes in *B. schroederi*, and comparative genomics revealed the potential common ability of host shift among ascaridoid parasites and the coevolution of these species. During its parasitism processes, *B. schroederi* appears to have evolved a thicker cuticle against the harsh intestinal environment and specialised its metabolism pathways to better utilize the limited nutrients observed during parasitism in the giant panda gut. Moreover, the enriched proteases in *B. schroederi* are linked to potential roles in host evasion and immunoregulation. These findings provide a useful resource that can be used in a wide range of fundamental biological studies of *Baylisascaris* and will strengthen the development of new interventions (drugs and vaccines) against baylisascariasis in the giant panda, which might constitute an epitome of wildlife conservation.

## Results

### Genome and gene sets

Using a combination of Illumina whole-genome shotgun technology, the PacBio single-molecule real-time (SMRT) sequencing platform and Hi-C scaffolding (Table S1), we produced a high-quality chromosome-level reference genome of *B. schroederi* that consisted of 293 megabases (Mb), had a scaffold N50 size of 11.8 Mb and was anchored to 27 chromosome-level pseudomolecules, which were numbered according to their collinearity with *A. suum* (Figure 1, Table 1, Figures S2 and S3, Table S2). The assembly size was larger than that of the horse *P. univalens* (253 Mb) [9], comparable to that of the swine *A. suum* (298 Mb) [10] and smaller than that of the dog *T. canis* (317 Mb) [11]. The GC content of the assembly was 37.59%, which is similar to that of *A. suum* (37.8%) but slightly lower than those of the *P. univalens* (39.1%) and *T. canis* (39.9%) assemblies. The completeness of the *B. schroederi* genome was estimated to achieve 97.86% (961/982) coverage using the core BUSCO genes [12] and 91.18% mapping using the transcriptomic data, which indicated that the assembly represents a substantial proportion of the entire genome (Table 1, Tables S3 and S4). The *B. schroederi* genome contained 12.02% repetitive sequences, which was equal to 35.3 Mb of the assembly, and these included 0.49% DNA transposons, 2.86% retrotransposons, 5.75% unclassified dispersed elements and 2.61% simple repeats (Table S5). Moreover, 6,190 transfer RNA (tRNA) and 978 ribosomal RNA (rRNA) genes were identified in the assembled genome, and the copy numbers reflected their codon usage in protein-coding regions (Figure S4, Tables S6 and S7, File S1).

**Figure 1.**
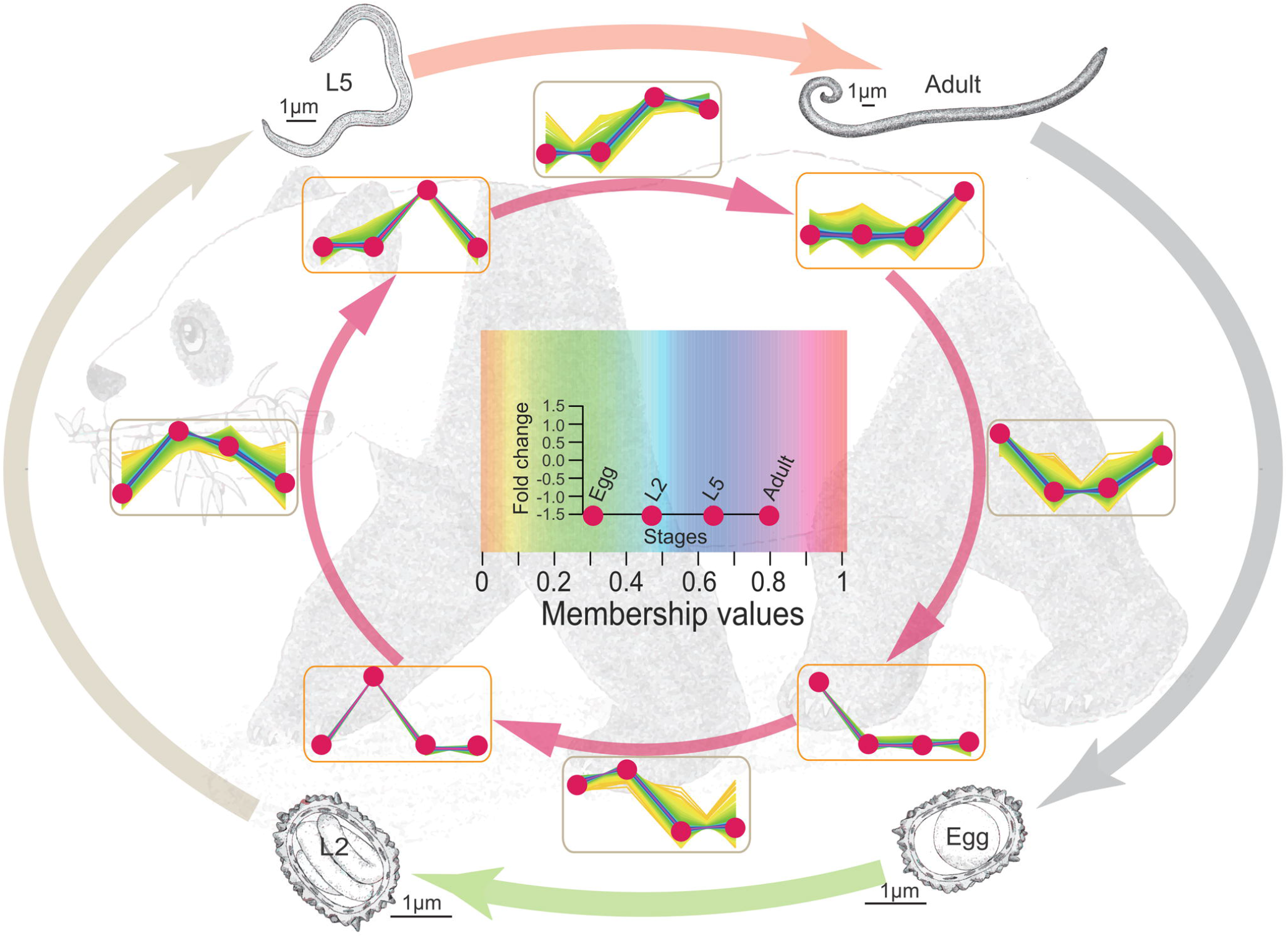
Genomic features of the *Baylisascaris schroederi* genome. The rings depict the following information with a window size of 100 kb: (a) Illumina sequencing coverage, (b) PacBio sequencing coverage, (c) repeat density, (d) gene density, (e) GC content, and (f) to (i) gene expression levels at the egg, L2, L5 and adult stages of *B. schroederi*, respectively.

**Table 1.** Statistics of the genome features of four ascaridoid species.

|  | *B. schroederi* | *A. suum* | *P. univalens* | *T. canis* |
| --- | --- | --- | --- | --- |
| **Versions** | BSFv2.0 | ASM18702v3 | ASM225920v1 | ADULT_r1.0 |
| **Genome size (bp)** | 293,522,654 | 298,028,455 | 253,353,821 | 317,115,901 |
| **Scaffolds** | 150 | 415 | 1,274 | 22,857 |
| **N50 (bp)** | 11,819,000 | 4,646,302 | 1,825,986 | 375,067 |
| **GC content (%)** | 37.59 | 37.8 | 39.1 | 39.9 |
| **Repeat (%)** | 12.02 | 11.1 | 7.7 | 12.9 |
| **Number of coding genes** | 16,072 | 16,778 | 14,325 | 18,596 |
| **Gene density (gene/Mb)** | 54.7 | 56.3 | 56.5 | 58.6 |
| **Mean protein length (aa)** | 492 | 436 | 470 | 385 |
| **BUSCO (complete/fragment, %)** | 92.2/5.7 | 89.1/7.2 | 91.0/5.8 | 87.0/8.0 |

*De novo* predictions, homology-based searching and deep transcriptome sequencing at multiple lifecycle stages of *B. schroederi* identified a total of 16,072 protein-coding genes, with an average length of 9,452 bp, an exon length of 155 bp and an estimated 9.5 exons per gene (Table S8), which were comparable to the data obtained for *A. suum, P. univalens* and *T. canis* [9–11]. In addition, 90.61% of the gene set was supported by the mapping of RNA-seq reads (Table S9) and had a homologue (BLASTP cutoff≤10-5) in *A. suum* (n=13,727; 85.41%), *P. univalens* (n=13,490; 83.93%) or *T. canis* (n=13,873; 86.32%); in addition, 11,135 (69.28%) were homologous among the four ascaridoid species, and 1,283 (7.9%) were ‘unique’ to *B. schroederi* because no homologs were detected in any other ascaridoids for which genomic data are currently available (Figure S5). Using this gene set, we then performed functional annotation with public databases. In total, 14,968 (93.13%) and 14,831 (92.28%) genes had homologues in the Nr and InterPro databases, respectively, whereas 10,331 (64.28%) and 4,414 (27.46%) genes contained Pfam domains and at least one transmembrane domain, respectively (Table S10). Notably, 3,465 (21.56%) genes of the 16,072 genes of *B. schroederi* had an ortholog linked to one or more of 134 known biological pathways (KEGG), and most of these genes can also be mapped to those in *C. elegans* (Table S11). Moreover, 28 genes were assigned to four groups of antimicrobial effectors, namely, cecropins, saposins, neuropeptide-like proteins (NLPs) and nematode antimicrobial peptides (AMPs) (Table S12, File S1).

### Genome evolution and host shift of ascaridoids

To determine the evolution of ascaridoid parasites in the context of nematodes, we inferred the phylogeny from 329 single-copy core orthologs across 12 nematode genomes using the maximum likelihood method (Figure 2A, Figure S6). Based on the phylogeny, the orders Ascaridida, Spirurina and Rhabditina were each treated as a monophyletic group in the phylum Nematoda, in accordance with the previously proposed molecular phylogeny [13, 14]. We further focused on the ascaridoid species in Ascaridida and found that the giant panda *B. schroederi* was more closely related to the pig *A. suum* and the horse *P. univalens* than to the dog *T. canis*, and these findings supported the hypothesis that *B. schroederi, P. univalens* and *A. suum* belong to the same family (Ascarididae), whereas *T. canis* belongs to the family Toxocaridae [15–17]. Building on the phylogeny, we estimated the divergence time of the ascaridoid parasites and other nematodes. Combining the previously published divergence dates with the fossil record [18–21], we estimated that the species of Ascaridida and Spirurina separated at 238 Mya and that the species of Ascaridida/Spirurina and Rhabditina diverged at 365 Mya (Figure 2A, Figure S7). Furthermore, among the order Ascaridida, the giant panda (*Baylisascaris*) and pig/horse (*Ascaris*/*Parascaris*) ascaridoids separated at 22 Mya in the early Miocene period, and the giant panda (*Baylisascaris*) and dog (*Toxocara*) ascaridoids diverged at 59 Mya in the late Paleocene period. Both of these divergence times appeared to postdate splits between *Baylisascaris* and *Ascaris*/*Parascaris* (~70 Mya) and between *Baylisascaris* and *Toxocara* (~200 Mya), respectively, that were previously estimated based on partial nuclear and mitochondrial genes [22] but agreed with an earlier speculation in which species of *Ascaris* and *Caenorhabditis* diverged at ~400 Mya [23].

**Figure 2.**
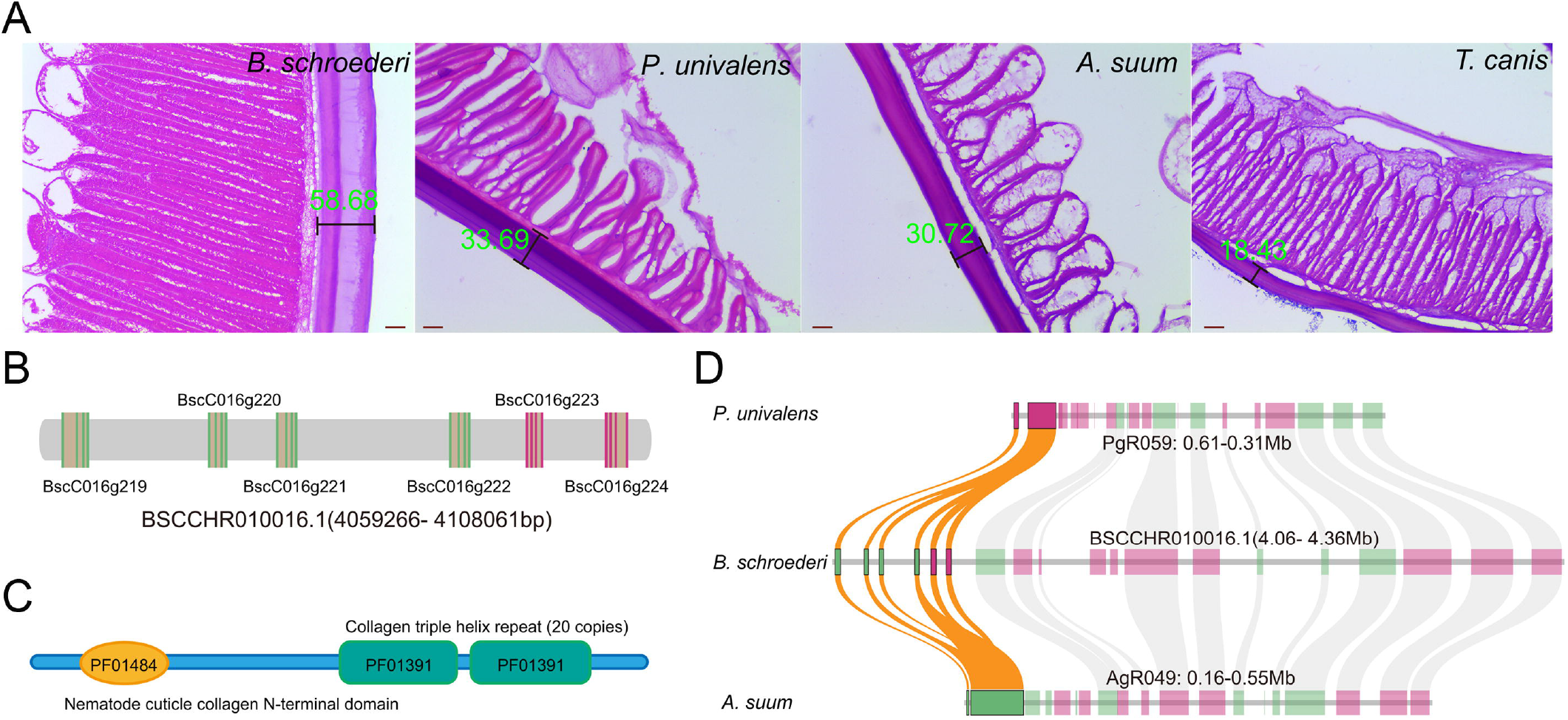
Phylogeny and inferred host shifts of *B. schroederi* and related ascaridoid species. **A**. The phylogenetic tree and divergence times of four ascaridoids and their hosts. The above tree of shows the relationship of the ascaridoid hosts include pigs, horses, giant pandas and dogs, which is estimated based on previous studies [22, 24]; the bottom tree shows the relationship of the ascaridoids included in this study, which is inferred based on 329 single-copy orthologs using RAxML and BEAST2. The color branches in the bottom tree indicate different ascaridoid species. The circles indicate species differentiation and the pentagrams denote host shifts and speciation. The dashed lines, circles, and pentagrams float on the host tree are transcribed from the ascaridoid tree. The arrows and corresponding lines indicate the direction of host shifts. **B**. Ks distribution of homologous genes among the four ascaridoids. **C**. GO enrichment of differentiation genes, which are mainly enriched in transferase, transporter, metalloendopeptidase, transmembrane transporter and ion channel.

Moreover, an assumed ‘host shift’ theory was developed based on the following results from our analysis of the divergence times and topological relationships among the four ascaridoid species and their host animals: (1) the divergence time (59 Mya) of *T. canis* and *B. schroederi* is close to that of the host dog and giant panda (61 Mya) [24], and the common ancestor of the giant panda ascaridoid and those of the pig and horse split from the ancestor of the dog ascaridoid following the differentiation of their own hosts; (2) *P. univalens* shared a lower similarity with *T. canis* than with *B. schroederi* and *A. suum*; (3) the three ascaridoid species (*B. schroederi, A. suum* and *P. univalens*) postdate the differentiation of their own hosts, and the divergence times between the giant panda and horse and between the pig and horse were 89.6 Mya [24] and 105.7 Mya [25], respectively; (4) the pig *A. suum* is closer to the horse *P. univalens* than to *B. schroederi*; and (5) the similarity of orthologous genes between *B. schroederi* and *A. suum* is higher (Wilcoxon signed rank test, *p*<2.2e-16) than that of orthologous genes between *B. schroederi* and *P. univalens* (Figure S8). We thus concluded that the common ancestor of the ascaridoids from the giant panda, pig and horse diverged from the ancestor of the dog ascaridoid as the dog and bear ancestors diverged. The ancestor of these three ascaridoids first colonized the panda ancestor; subsequently, the pig ancestor acquired the ascaridoid from panda ancestors via predation or food webs and fixed this parasitism until formation of its own ascaridoid *A. suum*; and the horse ancestor then acquired the ascaridoid from pig ancestors and gave rise to the horse ascaridoid *P. univalens* (Figure 2A). Furthermore, our analysis of the *Ks* distribution of orthologous genes between the four ascaridoid species also supported the ‘host shift’ assumption because the peak of the *Ks* distribution of orthologous genes between *B. schroederi* and *A. suum* was significantly skewed from that between *B. schroederi* and *P. univalens* (Figure 2B), which is in contrast to the theoretical observation that *A. suum* and *P. univalens* grouped in one branch and either between *A. suum* and *B. schroederi* or between *P. univalens* and *B. schroederi* should exhibit an overlapping peak in the *Ks* distribution. Furthermore, the peak value of the *Ks* distribution between *B. schroederi* and *A. suum* was lower than that found in the *Ks* distribution between *B. schroederi* and *P. univalens*.

In addition, based on pairwise comparisons of diverged genes retrieved from orthologous genes between *B. schroederi, A. suum* and *P. univalens*, we found that some diverged genes were shared among these three ascaridoids (Table S13A). Based on the consideration that host adaption-related genes would change before and after host shifts, these shared diverged genes likely provided us with opportunities to explore gene clues that underline host shifts. Analyses combining GO enrichment and functional annotations showed that most of the genes were enriched in ion channel activity (GO:0005216, *p*=2.60E-05), transporter activity (GO:0005215, *p*=3.54E-03), metalloendopeptidase activity (GO:0004222, *p*=7.71E-03), and transferase activity/transferring hexosyl groups (GO:0016758, *p*=2.43E-02) (Figure 2C, Table S13B), which play roles as material transport-related carriers, including sugar (and other) transporters, transmembrane amino acid transporter proteins, ABC transporters and ammonium transporter/ion transport proteins, and have functions in material metabolism, including glycolysis/gluconeogenesis, biosynthesis of amino acids, amino sugar and nucleotide sugar metabolism, glycerophospholipid metabolism and purine/pyrimidine metabolism (Table S14). These findings at least partly agree with the conclusion that the host shifts accelerate the divergence of these orthologs among these three ascaridoids to allow better adaptations to their new nutritional environment due to differences in host feedings. In addition, several genes, including those that encode immunoglobulins, lectins, flavin-containing amine oxidoreductases, thioredoxins, serpins and tetraspanins, were predicted to be involved in host-parasite immune interactions. For instance, flavin-containing amine oxidoreductases might modulate the levels of host amines (e.g., histamine) and trigger tissue damage in nematode infections [26, 27]. Tetraspanins bind the Fc domain of immunoglobulin (Ig) G antibodies and might help the parasites evade host immune recognition and complement activation [26, 28].

### Specialized nutrition

The long coevolutionary history between parasitic ascaridoids and their hosts has resulted in the relatively good tolerance of parasites in the host gut [22, 29, 30]. Food consumed by the giant panda is almost exclusively composed of bamboo. Thus, to survive in an environment where nutrients are relatively scarce and simple, *B. schroederi* might strengthen genes related to basal energy expenditure to meet its own nutrient requirements and the metabolism of major nutrients. Comparative genomic analysis showed that genes that encode transporters were under expansion and/or positive selection. The ABC transporter (PF00005), sugar (and other) transporter (PF00083), major facilitator superfamily (PF07690), MFS/sugar transport protein (PF13347), neutral and basic amino acid transport protein (solute carrier family 3, PF16028), transmembrane amino acid transporter protein (PF01490), excitatory amino acid transporter (sodium:dicarboxylate symporter family, PF00375) and long-chain fatty acid transport protein (AMP-binding enzyme, PF00501) gene families are expanded, and the former two are also identified with positive selection (Figure 3, Tables S15 and S16), which suggests an increasing ability to transport sugars, amino acids and fatty acids.

**Figure 3.**
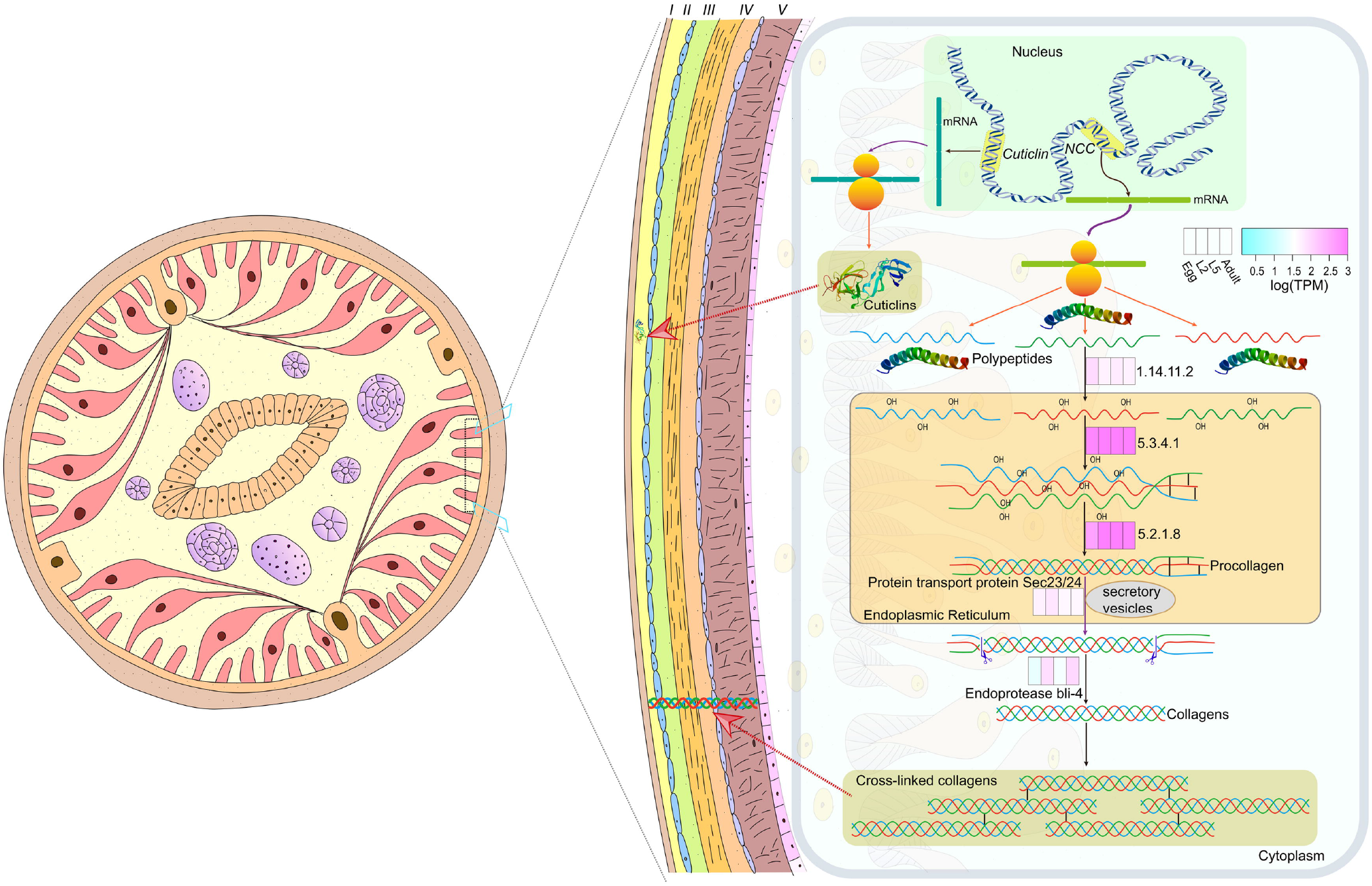
Nutrient transport and metabolism of *B. schroederi*. *B. schroederi* absorbs nutrients such as sugars, amino acids and fatty acids from the gut of giant pandas through transport proteins to enter cells and carry out metabolic processes in the cells. Different colored lines indicate different metabolic pathways: green, sugar metabolism; red, TCA cycle; blue, amino acid synthesis; dark brown, fatty acid synthesis; brown, glycogen synthesis; and pink, chitin metabolism. The dots indicate metabolic substrates or products. The number indicates the EC number of the enzyme, the double arrow shows that the enzyme-encoding gene that catalyzes the indicated step is expanded, and the asterisk means that the enzyme-encoding gene was under positive selection. The four colored boxes indicate the expression levels of the enzyme-encoding genes during the four developmental stages including eggs, L2s, L5s, and adult females.

In sugar metabolism, increased sugar transport capacity enhances sugar metabolism-related pathways to provide energy and increase intermediate products to improve the efficiency of conversion of other nutrients (e.g., amino acids and fatty acids). In the glycolysis/gluconeogenesis and citrate cycle (TCA cycle), the hexokinase (EC:2.7.1.1) pyruvate dehydrogenase E2 component (dihydrolipoamide acetyltransferase) (EC:2.3.1.12), citrate synthase (EC:2.3.3.1) and succinyl-CoA synthetase alpha subunit (EC:6.2.1.4)-related gene families were expanded (Fig. 3 and table S15). Moreover, the glyceraldehyde 3-phosphate dehydrogenase (EC:1.2.1.12), isocitrate dehydrogenase (EC:1.1.1.42), succinate dehydrogenase (ubiquinone) iron-sulfur subunit (EC:1.3.5.1), and related gene families were identified with positive selection (Table S16). Interestingly, the phosphoglycerate kinase (EC:2.7.2.3)-, pyruvate dehydrogenase E1 component alpha subunit (EC:1.2.4.1)-, and aconitate hydratase (EC:4.2.1.3)-related gene families were both under expansion and positive selection. These results showed that enhancing the activity of enzymes related to glucose metabolism promotes the formation of intermediate products and the conversion of other nutrients.

In addition, important enzymes for amino acid and fatty acid biosynthesis were also expanded, which resulted in the enhancement of their synthesis ability. In the biosynthesis of amino acids, the glutamate dehydrogenase (NAD(P)+) (EC:1.4.1.3)-, D-3-phosphoglycerate dehydrogenase/2-oxoglutarate reductase (EC:1.1.1.95)-, pyrroline-5-carboxylate reductase (EC:1.5.1.2)-, branched-chain amino acid aminotransferase (EC:2.6.1.42)-, phosphoglycerate kinase (EC:2.7.2.3)-, aconitate hydratase (EC:4.2.1.3)-and asparagine synthase (glutamine-hydrolyzing) (EC:6.3.5.4)-related gene families were expanded (Fig. 3 and table S15). The glyceraldehyde 3-phosphate dehydrogenase (EC:1.2.1.12)-and branched-chain amino acid aminotransferase (EC:2.6.1.42)-related gene families were under positive selection (Table S16). Interestingly, the branched-chain amino acid aminotransferase (EC:2.6.1.42)-related gene family was also under expansion and positive selection. Simultaneously, similar metabolic processes occur in fatty acid biosynthesis. The long-chain acyl-CoA synthetase (EC:6.2.1.3)-related gene family was expanded, and the mitochondrial enoyl-(acyl-carrier protein) reductase/trans-2-enoyl-CoA reductase (EC:1.3.1.38)-related gene family was positively selected (Figure 3, Table S16). All these metabolic processes indicate that *B. schroederi* can use host nutrients to synthesize nutrients to compensate for inadequate nutrition in the giant panda gut. The transcriptome analysis showed that these genes are highly expressed in the intestine of worms, which further proves that the worms replenish nutrition by enhancing metabolic activities to adapt well to the nutritional environment of the giant panda gut.

### Specialized cuticle in *B. schroederi*

The nematode cuticle plays a protective role against a variety of external biotic and abiotic stresses and is composed of five layers, including the surface coat, epicuticle and cortical, medial and basal layers [31–34]. Although different layers contain distinct molecular assemblies, the cuticle is mainly formed from collagens, cuticlins, chitin and small amounts of lipids [34–36]. Collagens, which are the structural proteins in cuticles and comprise the major component of the extracellular matrix, are synthesized through a multistep process that includes prolyl 4-hydroxylation, procollagen registration and trimerization, transport from the endoplasmic reticulum, and procollagen processing and cross-linking, and more than 170 genes are involved in the production of this protein, similar to the phenomenon in vertebrates [34–38]. In addition, cuticlins, which are another structural component of the cuticle and are abundant in the outermost cortical layers, are hypothesized to be enzymatically polymerized to constrict the seam cell-derived cuticle and thereby form the distinctive cuticular alae in *C. elegans* [33, 34, 39].

Under this context, we retrieved the genes encoding cuticle collagens from the *B. schroederi* gene set. We identified 158 gene copies, and each expressed product contained a nematode cuticle collagen N-terminal domain and/or collagen triple helix repeats (n=20) (Figure 4, Table S17). A transcriptome analysis showed significantly differential expression of the genes encoding cuticle collagens during the development of *B. schroederi*, and the genes presented quite high expression levels at the L5 and adult stages, which indicated that a large number of collagens are needed for formation of the worm cuticle. Collagen synthesis is catalyzed by prolyl 4-hydroxylase (EC:1.14.11.2), protein disulfide-isomerase (EC:5.3.4.1) and peptidyl-prolyl cis-trans isomerase A (EC:5.2.1.8), and this step is followed by cleavage at the N-and C-termini by endoprotease and zinc metalloproteinase and then maturation and structural cross-linking by dual oxidase (EC:1.6.3.1). These enzymes, which the exception of zinc metalloproteinase and dual oxidase, are highly expressed at the L5 and adult stages and contribute to the formation of a thick exoskeleton for body protection against threats posed by the complex intestinal environment of the host giant panda, particularly its special bamboo-dominated diet habit. In addition, 32 cuticlins (PF00100, zona pellucida-like domain) were also identified in *B. schroederi* (Figure 4, Table S17).

**Figure 4.**
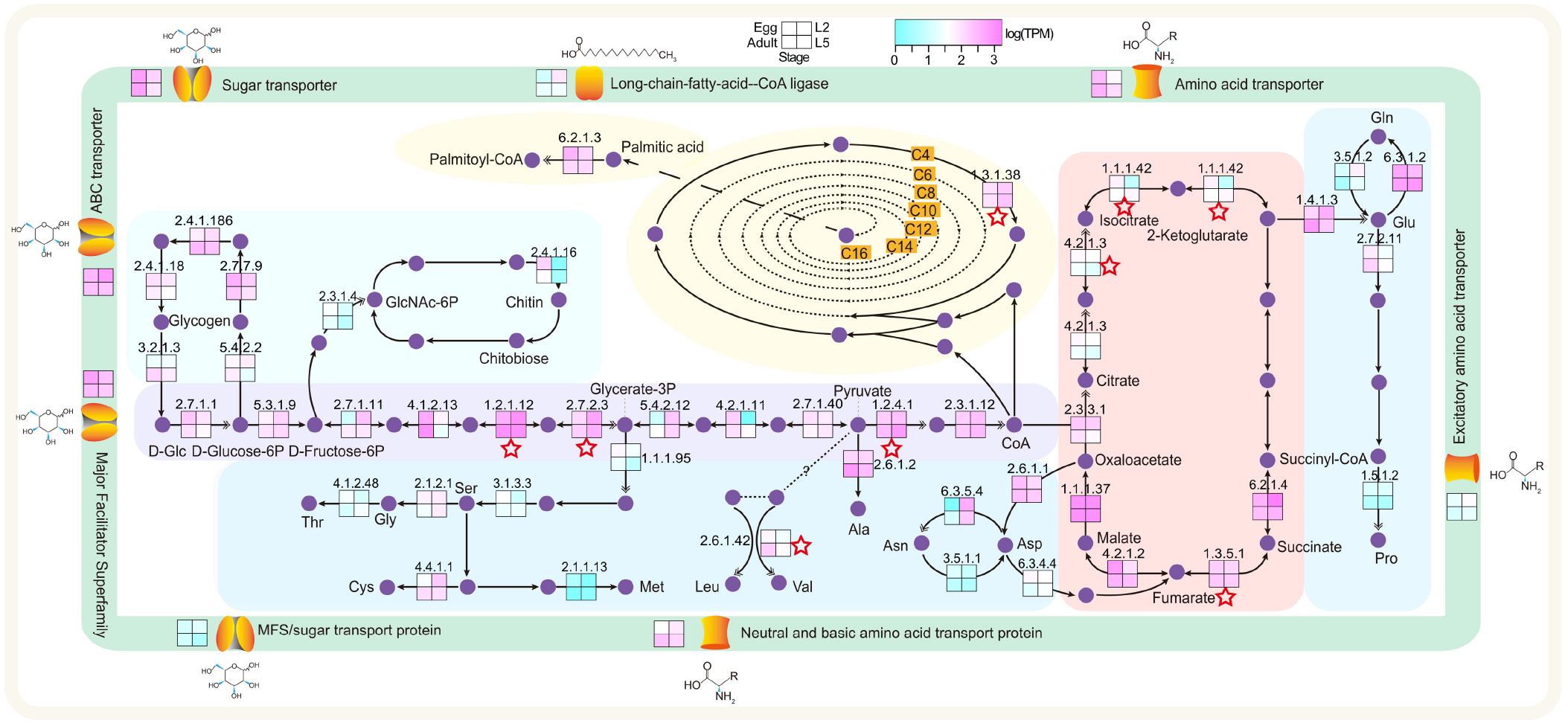
Composition of and related gene expression in the *B. schroederi* cuticle. Cuticle of *B. schroederi* is magnified from its transverse section model to illustrate its structure that includes the surface coat (I), epicuticle (II) and cortical (III), medial (VI) and basal layers (V) and composition that is made up of collagens, cuticlins, chitin and lipids. Transcriptome analysis showed significantly differential expression of the genes involved in biosynthesis of cuticle collagens and cuticlins during the development of *B. schroederi*. Within the biosynthesis of cuticlins and collagens, cuticlins are enzymatically polymerized to constrict the seam cell-derived cuticle and form the distinctive cuticular alae, which are predominant in the outermost cortical layers; while collagens are synthesized through a multistep process that includes prolyl 4-hydroxylation, procollagen registration and trimerization, transport from the endoplasmic reticulum, and procollagen processing and cross-linking, which results in construction of the major component of the extracellular matrix of the epicuticle and cortical, medial and basal layers of the nematode cuticle. For expression levels of genes related to biosynthesis of collagens, the mean centered log-fold change in expression is plotted for each of three biological replicates at each life stage in the following order: eggs, L2s, L5s, and adult females. All genes in each cluster are drawn with different colors. The red and blue bars indicate low and high deviation from the consensus profile, respectively.

Interestingly, our speculation was confirmed by histological examinations (Figure 5A, Figure S9, Table S18), which revealed that the *B. schroederi* cuticle was significantly thickened compared with those of the other three ascaridoids included in this study, namely, *A. suum, P. univalens* and *T. canis* (*p*<0.01). To exclude whether this thickness difference is derived from species variations among ascaridoids, we also introduced the ursine *Baylisascaris transfuga* for comparisons because this ascaridoid is congeneric with *B. schroederi* in the genus *Baylisascaris* (Figure S10). Notably, *B. schroederi* has a markedly thicker cuticle than *B. transfuga* (*p*<0.01). We further compared the cuticle-related genes among *B. schroederi, A. suum, P. univalens* and *T. canis* (*B. transfuga* was not included because its genome has not yet been sequenced) and found that cuticle collagens were duplicated after the separation of *B. schroederi* and *Ascaris*/*Parascaris* (Figure 5A and C) and that these genes were highly expressed in *B. schroederi* at the L5 or adult stages (Table S16). A structural analysis showed that most of these cuticle collagens were present in collinear regions among these three ascaridoids in the form of tandem repeats with equal sequencing depths (Figure 5D, Figure S11), which suggests the authenticity of gene expansion rather than genome annotation errors. In addition, peptidyl-prolyl cis-trans isomerases (EC: 5.2.1.8) and cuticle collagens were positively selected. Such adaptive selections would undoubtedly enhance the functions of these genes in the cuticle of *B. schroederi*. Combined, these results suggest that through copy-number increases and the functional strengthening of genes involved in cuticle collagen formation, *B. schroederi* has evolved a thicker cuticle as armor to protect itself from the sharp-edged bamboo culm/branch-enriched intestinal environment during parasitism in the giant panda as it experienced a significant dietary change from meat to bamboo throughout history.

**Figure 5.**
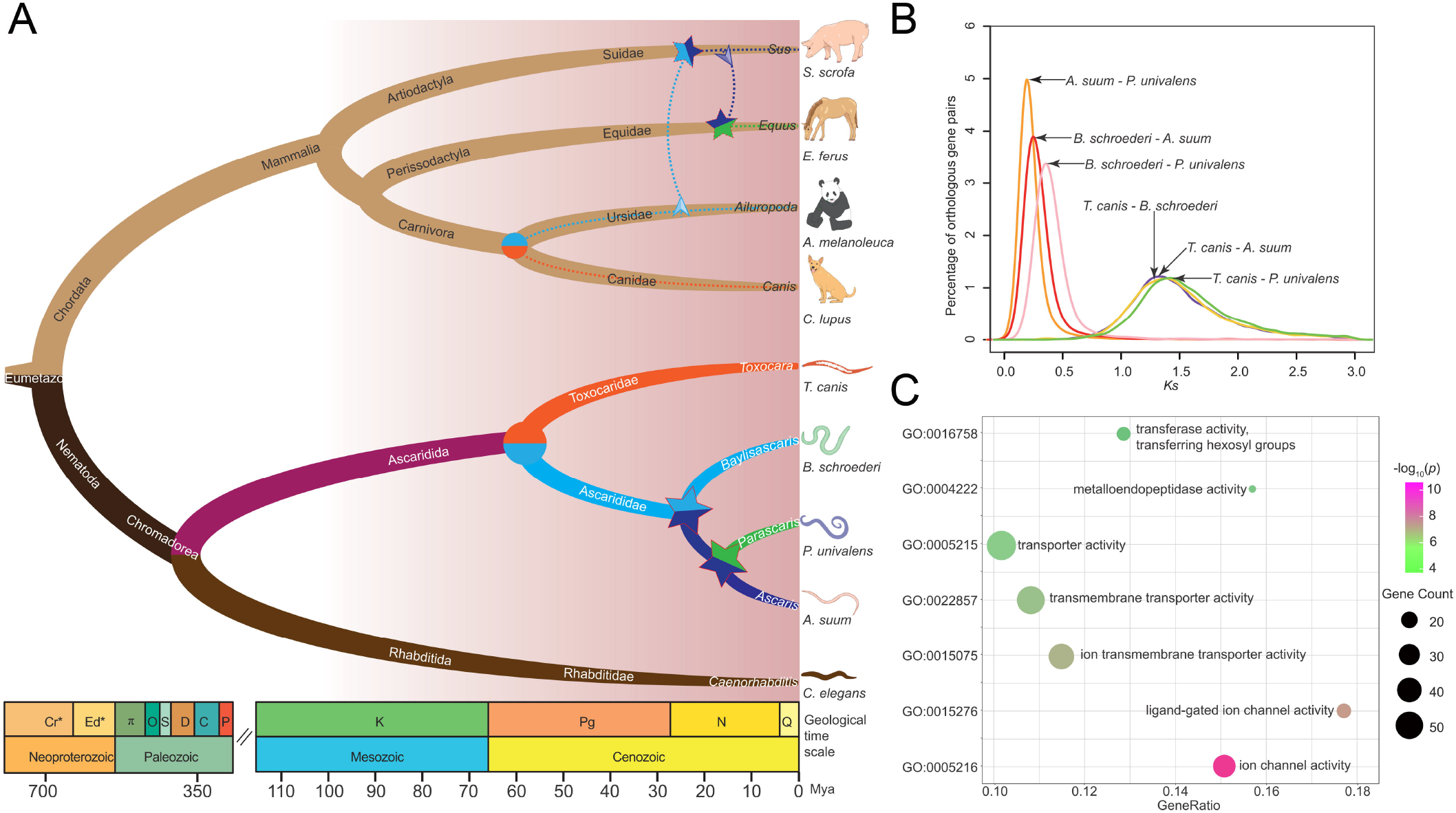
Cuticle thickness and expanded genes related to cuticle biosynthesis in ascaridoids. **A**. Cuticle thickness of *B. schroederi, A. suum, P. univalens* and *T. canis* under 400×. The scale bars denote 20 µm. **B**. The genes encoding cuticle collagens are presented in tandem on the genome. Green means that the genes are on the positive strand while red means that the genes are on the negative strand. **C**. The Pfam domain of cuticle collagens that includes one “nematode cuticle collagen N-terminal domain” and two “collagen triple helix repeat (20 copies)”. **D**. Genes encoding nematode cuticle collagens were tandemly duplicated in syntenic blocks between *B. schroederi* and *A. suum*/*P. univalens*. The highlight colors represent genes encoding nematode cuticle collagens, and the dimmed colors represent other genes in syntenic blocks.

### Transcriptomic changes in the life stages

To understand the developmental processes of *B. schroederi*, we profiled genes that were differentially transcribed among eggs, infective second-stage (L2) larvae, initially intestine-inhabiting fifth-stage (L5) larvae and adults across the lifecycle (Figure 6, Tables S19and S20). We found 14,178 genes that were significantly expressed during at least one stage, and 11,510 genes were differentially expressed among the four life stages. Furthermore, these 11,510 genes were grouped into expression clusters to uniquely describe each life stage and two life stages, and expression clusters showing a stepwise increase or decrease in expression corresponding to some transitions through the lifecycle were also included (Figure S12). The genes that were upregulated during development from egg to the infectious L2 stage included those involved in the chromatin assembly, cellular component organization and morphogenesis (Tables S19 and S20), which is in agreement with the progression from the embryonation to motile and infective larval stage. We simultaneously noted that the L2 stage was characterized by an increased number of transcribed genes related to signaling pathways, cell communication, response to stimulus, cellular homeostasis and molecular binding and/or transport (Tables S19 and S20), which mirrored the larval adaptation to external conditions and increased the need to detoxify build-up endogenous wastes, consistent with the results from previous studies in *T. canis* [11, 40]. In addition, the decrease in the transcription of genes associated with catalytic activity, oxidoreductase activity and electron transfer activity as well as different metabolic processes, including organic cyclic compound metabolic process, amino acid metabolic process and lipid metabolic process, observed at the L2 stage (Table S20) also supports the notion that the larvae experience a quiescent state that allows their adaptation to a reduced metabolic rate in order to survive for extended periods under outside conditions [41, 42].

**Figure 6.**
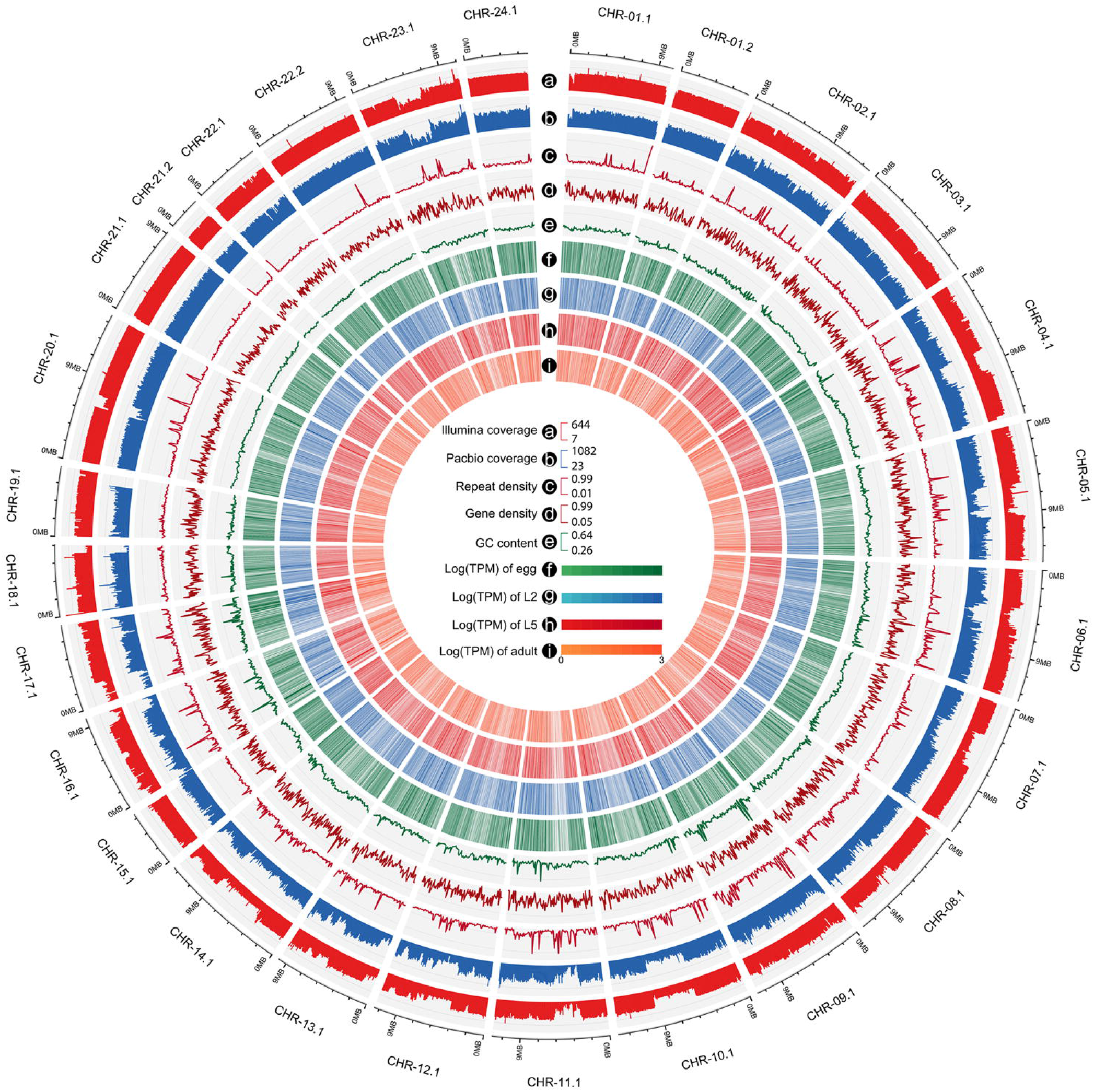
Example of differential gene expression clusters during the *B. schroederi* life cycle. The developmental transcriptomes of *B. schroederi* were sequenced in triplicate at four stages across the lifecycle (1st order): embryonated eggs (eggs); the second-stage larvae (L2s); the intestine-inhabited fifth-stage larvae (L5s); and adult females. A subset of the 11,510 differentially expressed genes (FDR<0.001, min fold=4) were grouped into expression clusters that describe the genes specifically upregulated at various life stages. Clusters that uniquely describe each life stage and describe two life stages are identified (2nd order). For all expression clusters, the mean centered log-fold change in expression is plotted for each of three biological replicates at each life stage in the following order: eggs, L2s, L5s, and adult females. All genes in each cluster are drawn with different colors. The red and blue bars indicate low and high deviation from the consensus profile, respectively.

L5 is the first intestine-inhabiting stage of *B. schroederi*, and its transition from the tissue/organ migrating larval stage was reflected by a significant upregulation of genes involved in metabolic processes, oxygen transport and the actin cytoskeleton as well as cuticle development (Table S19). The genes encoding protein kinases/kinases, peptidases, phosphatases, transferases and hydrolases, which are possibly associated with food degradation and digestion in *A. suum* and *T. canis* [10, 11], were also upregulated (Tables S19 and S20). We also noted significantly increased transcription of genes encoding enzymes related to cell redox homeostasis, including glutathione S-transferase, arylesterase, oxidoreductase and glutathione peroxidase and/or peroxiredoxin, which likely reflects the maintenance of the redox balance in response to the accumulation of the end products from anaerobic metabolism and the clearance of reactive oxygen species from endogenous metabolic activities during infection. In addition, the development process from L5 to adulthood was characterized by gene sets that were enriched in genes associated with metabolic processes, hormone mediated growth and development and embryogenesis in adult females (Tables S19 and S20). For instance, genes involved in amino sugar/carbohydrate metabolisms, insulin-like growth factor binding, steroid mediated growth and embryonic division were significantly upregulated, and this upregulation was accompanied by increased expression of genes involved in DNA replication/repair during this transition.

### Insights into new interventions

Because the current excessive use of a small number of drug classes for the treatment of baylisascariasis in the captive panda population has resulted in the emergence of drug resistance, the *B. schroederi* genome sequence theoretically provides an alternative approach to drug target discovery and repurposing [43]. We identified 1,093 essential genes linked to lethal-gene-knockdown phenotypes in *C. elegans*, and 454 of these were shared with the ChEMBL database (Table S21). One hundred ninety-four of these 454 genes were deemed one-copy orthologs and were absent in hosts (Table S21). Thus, we focused on this gene set and gave the highest priority to genes that are inferred to be highly expressed and to function as enzymatic chokepoints [42, 44, 45]. Under such strict criteria, druggable candidates, including peptidases (n=3), serine/threonine kinase (n=1) and protein phosphatases (n=2) (table S21), represent proven targets of norcantharidin analogs with nematocidal activity [46, 47]. The peptidases were threonine, serine and metalloenzymes, whereas the protein phosphatases consist of Ser/Thr phosphatases. The transporters are also validated targets for many current antihelminthtics, including imidazothiazole derivatives (including levamisole), macrocyclic lactones, cyclic depsipeptides and AADs [42, 48–50]. We identified four transporters in the *B. schroederi* genome (Table S21). The combined list of prioritized targets of drug candidates could prompt the rational design of anthelmintics, particularly when these targets exert antinematodal effects *in vitro*, as demonstrated through larval development assays, and *in vivo* in the giant panda.

In addition to drug target discovery, vaccine candidates that should be both immunologically accessible and crucial for parasite survival were also mined from the *B. schroederi* genome. Excretory/secretory (E/S) proteases and protease inhibitors appear to meet the requirements because they are secreted into the host, are thus exposed to the host’s immune system and can modulate the immune system of the host to promote parasitism. We surveyed genes encoding E/S proteins (Table S22, File S1), particularly proteases and protease inhibitors, that were expressed during the parasitic stages of *B. schroederi* and were parasite-specific genes (no orthologs in host mammals). Such screening yielded 85 proteases, which mainly included protease inhibitors (n=61); among these vaccine candidates, cysteine peptidases and thioredoxins were expressed at the L5 and adult stages (Table S23). We also noted the substantial diversity among the cysteine peptidases that contained cathepsin L and cathepsin W, and most of these, such as cathepsin L, have been under close scrutiny as vaccine candidates [51, 52]. Moreover, other E/S proteases, including serine peptidases and serpins that were upregulated at the parasitic stages, have also been observed, although their feasibility for the development of vaccines remains under evaluation [53, 54]. Therefore, a combination of genomic data and animal experimentation should advance the screening and development of vaccines in the future.

## Discussion

The diet of the giant panda, which is an endangered, herbivorous species, is made up almost exclusively of low-protein and high-fiber bamboos [1, 55, 56]. This high degree of specialization in low-quality foods not only shapes the panda’s behaviors, allowing it to cope with food challenges involving the levels and balance of essential nutrients, but also renders the adaptation of its tract-inhabiting organisms, including parasitic nematodes, to the harsh (sharp-edged bamboo culm/branch-enriched) intestinal environments of the pandas. The nematode *B. schroederi* is the only endoparasite that appears to be consistently found in pandas and is the leading cause of death from primary and secondary infections in wild pandas [4, 5]. Increased lines of evidence show that *B. schroederi* can grow to a body size comparable to those of other ascaridoid parasites, including *A. suum* in pigs and *P. univalens* in horses [4, 8], which suggests that this parasite has highly evolved to adapt to its host. Given the long coevolution between parasites and their hosts, it would be intriguing to explore the speciation of *B. schroederi* and its parasitic adaptation to the unique gut environment of the panda and to seek alternative measures for the prevention and control of infections. In this study, we decoded the genome of *B. schroederi* and found molecular clues related to host shift to illustrate the speciation and molecular evidence of the cuticle thickness and thus explain gut adaptation. We also characterized the key molecules involved in development or host–parasite interactions and their potential as intervention targets for *B. schroederi*. These results provide new insights into the biology and evolution of *B. schroederi* and contribute to the future development of novel treatments for baylisascariasis in pandas.

According to the genome-wide phylogeny, we inferred that among the order Ascaridida, *B. schroederi* is closer to *A. suum* and *P. univalens* than to *T. canis* and that *A. suum* shares the closest relationship with *P. univalens*. Based on the available fossil evidence, we further estimated that the divergence between *B. schroederi* and *T. canis* occurred markedly earlier than the separation from *A. suum* and *P. univalens* (59 vs 22 Mya). However, a comparative genomics analysis revealed that the similarity of orthologous genes from *B. schroederi* and *A. suum* is higher than that of orthologous genes from *B. schroederi* and *P. univalens*, and *P. univalens* shares the lowest similarity to *T. canis*. Considering morphometric and distribution data of these four ascaridoids as well as the historical biogeography and phylogeny of their own hosts (i.e., dogs (*T. canis*), pandas (*B. schroederi*), pigs (*A. suum*) and horses (*P. univalens*)) [15, 16, 22, 24], two host-shifting events likely occurred after divergence of the common ancestor of ascaridoids between dogs and pandas. In addition, the occurrence of *A. suum* appears consistent with speciation following a host colonization event from pandas to pigs apparently from a carnivoran source in sympatry, and the occurrence of *P. univalens* appears consistent with speciation following a host colonization event from dogs to horses apparently from predation or food webs. Such history of host colonization is compatible with the current tree topology and coincides with historical evidence of the spatiotemporal coappearance of the panda ancestor primal panda *Ailurarctos lufengensis* and the pig ancestor Eurasian wild boar *Sus scrofa* in the late Miocene and Pliocene and a recent molecular inference of a wide host-shifting origin for ascaridoid nematodes [22]. Nevertheless, the timing and geographic source for these ascaridoids cannot be elucidated in detail based on the currently available data and the reduced and relictual distribution of giant pandas. Future parasitological inventory among a wider host range in a region of sympatry is necessary to demonstrate that each ascaridoid species has a narrow host range and might now be limited to the present host [57]. In addition, the apparent differences among the genomes of *T. canis* in dogs, *B. schroederi* in pandas, *A. suum* in pigs and *P. univalens* in horses, coupled with their divergent biogeographic and ecological histories, also suggest that this system is a good model for exploring the complexities of diversification and faunal assembly in the evolution of the host range and the associations among ascaridoids (e.g., refs. [58–61])

In general, the nematode cuticle is an extremely flexible and resilient exoskeleton and plays vital roles against external stresses. This exoskeleton is composed primarily of cross-linked collagens, cuticlins, chitin and small amounts of lipids [34–36, 62]. In *B. schroederi*, we observed an accelerated evolution of genes related to cuticle biosynthesis, including the significantly higher expression of genes encoding the nematode cuticle collagens, chitin synthase, DP-N-acetylglucosamine pyrophosphorylase, and peritrophin-A chitin-binding proteins at the adult stage compared with that at other stages across the development of this parasite, and the significant expansion of genes such as collagens and peritrophin-A chitin-binding proteins compared with those in other ascaridoids, namely, *T. canis, A. suum* and *P. univalens* [9–11]. This overexpression and expansion of genes responsible for cuticle formation in *B. schroederi* suggest its parasitic adaptation to the intestinal environment of the panda, which is fully filled with highly fibrous and sharp-edged components of bamboos. This conclusion is further supported by our analysis of positive selection, which revealed that genes encoding cuticle collagens, peritrophin-A chitin-binding proteins, GlcN6P synthase and UDP-N-acetylgalactosamine diphosphorylase are also positively selected in *B. schroederi* compared with those in *A. suum, P. univalens* and *T. canis*. This molecular evidence, together with morphological and anatomical observations among these ascaridoid species, including a congeneric *B. transfuga* from bears (Fig. 4), reinforces the assumption that *B. schroederi* might have evolved a thicker cuticle as an armor to protect itself from the harsh external environment during its gut parasitism in pandas. Because the panda retains the alimentary tract of bears but evolved into a bamboo-eating herbivore, unlike other members of Ursidae, which are carnivores or omnivores, further studies that include the bear *B. transfuga* for genome comparison might be needed to illustrate the significantly thickened cuticle that is only present in the panda *B. schroederi*.

In this study, we present a chromosome-level genome assembly of the giant panda roundworm *B. schroederi* and uncover an evolutionary trajectory accompanied by host-shift events in ascaridoid parasite lineages after host separations and an increased cuticle thickness and efficient utilization of host nutrients in *B. schroederi*, which guarantee its gut parasitism in giant pandas. We also found a broad range of key classes of molecules involved to host-parasite interplay and immunoregulation that could serve as potentially ideal targets by developing new and urgently needed interventions (drugs, vaccines and diagnostic methods) for the control of baylisascariasis. These genome resources not only enable the transition from ‘single-molecule’ research to global molecular discovery in *B. schroederi* but should also contribute to the protection of giant pandas by providing a novel treatment for baylisascariasis.

## Materials and methods

### Samples and preparations

Adult worms of *B. schroederi* were collected from naturally infected giant pandas at Chengdu Research Base of Giant Panda Breeding, Chengdu (Sichuan, China). Embryonated eggs were obtained 2 cm proximal to the uteri of the females. The second-stage larvae (L2s) were harvested using established *in vitro B. schroederi* egg culture protocols. Briefly, after filtering through a

100-µm nylon sieve filter, washing with PBS, and centrifugation, the egg suspension obtained from the uteri was placed into 100-mm culture dishes and maintained at ambient room temperature for at least 60 days to embryonate the eggs to an infective L2 stage. The eggs with well-formed and ensheathed larvae in the suspension were counted, and the suspension was then stored at 4 °C until use. L5s (n=25) were occasionally isolated from naturally infected captive giant pandas at the Chengdu Research Base of Giant Panda Breeding. These larvae together with adult worms were washed extensively in sterile physiological saline (37 °C), snap-frozen in liquid nitrogen and then stored at −80 °C until use. All the samples of other ascaridoid specimens, including *B. transfuga, A. suum, Parascaris univalens* and *T. canis*, were also derived from naturally infected polar bears, pigs, horses and dogs, respectively, and provided by the Department of Parasitology, College of Veterinary Medicine, Sichuan Agricultural University.

Genomic DNA was extracted from the freshly collected middle portion of *B. schroederi* to construct one paired-end library and one SMRT library. Messenger RNA was isolated from *B. schroederi* embryonated eggs, L2s, L5s and adult females for the construction of paired-end cDNA libraries (300 bp). To further verify the identity of the specimen, the ITS1 and ITS2 sequences of nuclear ribosomal DNA (rDNA) were amplified by PCR and compared with those previously reported for *B. schroederi* (Accession number: JN210912) [63].

### Genome survey analysis

To survey the characterization of the *B. schroederi* genome, a 300-bp pair-end library was constructed, and a total of ~10-Gb next-generation sequencing data were generated using the Illumina sequencing platform (Hiseq4000) (Table S1). Adaptor sequences, PCR duplicates and low-quality sequences were removed from the raw data to generate high-quality sequences. K-mer (17) statistics of the high-quality sequences were calculated by Jellyfish (version 2.1.3) [64] with “-C -m 17”. GenomeScope (version 2.0) [65] software was used to estimate the size, heterozygosity and repeat content of the *B. schroederi* genome (Figure S13).

### Genome sequencing and assembly

One cell run of single-molecule long reads was generated with the PacBio Sequel II platform (table S1). A total of 189-Gb long subreads (97,400,959,204 bases, ~332× based on the estimated genome size) were generated and *de novo* assembled using CANU (version 1.8) [66]. The parameters were optimized for heterozygotic genomes according to the authors’ documentation. The initial CANU assembly was corrected using a combination of long and short reads using Pilon (version 1.23) [67] with the default parameters. Duplicated assembled haploid contigs were purged using Purge Haplotigs (version 1.1.1) [68], which reduced the assembly from 559 Mb to 299 Mb. A Hi-C library was constructed with *Hind*III as the digestion enzyme and sequenced in two batches with the Illumina HiSeq4000 and NovaSeq platforms. The purged contigs were anchored into superscaffolds using the Juicer and 3d-dna pipelines. The generated assembly files were visualized and manually optimized using the built-in assembly tool (JBAT) of Juicebox. Twenty-one pseudomolecules were preliminarily generated with the 3d-dna pipeline. After breaking weak or ambiguous contact links between large TAD blocks and rebuilding the boundaries, a total of 27 pseudomolecules were generated. Note that these pseudomolecules did not represent complete chromosomes, and downstream synteny analysis with the *A. suum* genome showed that at least six molecules were likely partial chromosomes, which might reduce the number from 27 to 24. However, due to the lack of direct evidence of the karyotype of *B. schroederi*, we did not modify the result. Finally, the genome assembly contains 27 chromosome-level pseudomolecules and 123 unplaced scaffolds. The completeness of the assembly was assessed through Benchmarking Universal Single-copy Orthologs (BUSCO) analysis (Version 3.0.2, lineage dataset: nematoda_odb9) [69] and using RNA-seq data.

### Identification of repeat elements and noncoding RNAs

RepeatMasker (version 4.0.5, http://www.repeatmasker.org/) with the default parameters was applied to identify the dispersed repeats and tandem repeats. The species-specific repeat library was constructed with RepeatModeler (version 1.0.5, http://www.repeatmasker.org/). Using this library, repetitive sequences were further annotated and classified with RepeatMasker (http://www.repeatmasker.org/). The tRNA genes were predicted by tRNAscan-SE (version 1.3.1) [70] with general eukaryote parameters. The programs RNAmmer (version 1.2) and rfam_scan.pl (version 1.2) [71] were used to predict the large ribosomal subunit (LSU) and small ribosomal subunit (SSU) rRNA genes, respectively.

### Gene annotation

Protein-coding genes were annotated using a combination of *ab initio* gene prediction, homology-based gene prediction and transcriptome-based prediction. A total of 12 RNA-seq libraries were used to construct the transcripts by applying the HISAT2 (version 2.1.0) and StringTie (version 1.3.4) pipelines [72]. All constructed transcripts were combined using TACO (version 0.7.3) [73]. The ORFs on the transcripts were extracted with TransDecoder (version 5.5.0) [74]. The complete CDS from the TransDecoder result was used as the training set for *ab initio* prediction, which was performed with the BRAKER2 pipeline (version 2.1.5) [75]. All protein sequences of the previously sequenced nematode genomes were mapped to the genome using GenomeThreader (version 1.7.1) [76]. EVidenceModeler (version 1.1.1) [77] was employed to integrate the results from the three prediction methods and thus generate a consensus gene set, and the resulting set was further curated by removing frameshift and redundancy using the GFFRead (version 0.11.6) [78] tool from Cufflinks. Gene function annotation was performed using BLASTP (-evalue 1e-3) with public databases such as the nonredundant protein database (Nr) [79] and the KEGG database [80, 81]. InterproScan [82] was used to identify domains of the predicted proteins, assign GO terms to the predicted genes and classify the functional annotations.

### Gene family analysis

Protein sequences from *B. schroederi* and 11 other nematodes (*A. suum, P. univalens, T. canis, Loa loa, B. malayi, H. bacteriophora, H. contortus, P. pacificus, C. elegans, M. hapla* and *T. spiralis*) were used to analyze the gene family. Proteins with a length shorter than 30 aa or a frame shift were removed from the protein set, and the program OrthoFinder (version 2.3.3) [83] with the default parameters was used to construct the gene families and infer orthologous and paralogous genes. CAFE (version 3.0) [84] was utilized to identify the gene families that underwent expansion or contraction using the ultrametric tree inferred by BEAST2 and the estimated birth-death parameter λ.

### Homolog comparison

An all-to-all BLASTP (-evalue 1e-3 -outfmt 6) analysis of the proteins was performed to calculate the pairwise similarities. The *Ks* values of orthologous genes were calculated using codeml with the setting “runmodel = −2, CodonFreq = 2, model = 0, NSsites = 0”.

To compare the similarities among Ascarididae species at the gene level, a similarity analysis of orthologous genes was performed with the following steps:

(1) *A. suum* and *P. univalens* were selected as the query species to analyze which species is more similar to the target species *B. schroederi*; *B. schroederi, A. suum* and *P. univalens* were selected as the query species to analyze which species is more similar to the target species *T. canis*;

(2) The best-match orthologous genes of the target and query species were obtained from MCScan blocks;

(3) The similarity index (SI) was calculated using the formula SI=S/L, where S represents the alignment score of a pair of proteins and L represents the target protein length;

(4) Based on the SI value, the Wilcoxon signed rank test was used to analyze whether the similarity between different query species and target species was significantly different (p<=0.05, one-sided).

### Phylogeny construction

A total of 329 single-copy gene families were obtained, and the corresponding protein sequences were extracted. Individual protein alignment for each gene family was performed using Clustal Omega (version 1.2.1) [85] with the default settings, and gaps in the alignments were removed using the program trimAl (version 1.4) [86]. The alignments with a length of at least 100 aa were concatenated with a Perl script. The best amino acid substitution model for the protein alignment was estimated by ProtTest (version 3.4.2) [87] with the parameter “-IG -F -AIC -BIC -S 2 -all-distributions -tc 0.5”. The maximum likelihood tree was constructed using RAxML (version 8.0.24) [88] with the following parameters: 1) bootstrapping replicates, 200; 2) substitution model, LG+I+G+F; and 3) outgroup, *T. spiralis*.

### Divergence time estimations

The divergence time was estimated from the protein alignment by BEAST2 (version 2.5) [89] with the following parameters: 1) site Model, WAG+I+G+F; 2) clock model, relaxed clock log normal; 3) priors, calibrated Yule model; 4) time calibration: the split time (382-532 Mya) of Chromadorea and Enoplea [18], the split time (280-430 Mya) of *Pristionchus* and *Caenorhabditis* [19] and two fossil times (~396 and 240 Mya) for Enoplia [20] and Ascaridoidea [21], respectively; 5) chain length per MCMC run, 10,000,000. A consistent tree with divergence times was inferred by TreeAnnotator with a maximum clade credibility method and displayed using FigTree (https://github.com/rambaut/figtree/releases).

### Identification of positively selected genes

Orthologous genes of the four Ascarididae species (*B. schroederi, A. suum, P. univalens* and *T. canis*) were extracted from the OrthoFinder results to identify positively selected genes (PSGs). Multiple protein sequence alignments were performed with Clustal Omega and converted to corresponding CDS alignments using an in-house Perl script. Gaps in the CDS alignments were removed with the program trimAl. The Codeml program with modified branch-site model A (model = 2, NSsites = 2) as implemented in the PAML package (version 4.9) [90] was used to identify PSGs. The alternative hypothesis with estimated ω2 (fix_omega=0 and initial omega=1.5) and the corresponding null model with fixed ω2=1 (fix_omega=1 and omega=1) for the lineage *B. schroederi* (foreground branch) were used to calculate the omega values and log likelihood values, respectively. The likelihood ratio test (LRT) for selection of the lineage of *B. schroederi* was performed based on the likelihood values obtained from the two models. Genes with p<=0.05 were PSGs.

### Transcriptome analysis

Adaptor sequences, contaminants and low-quality sequences were removed from the raw RNA-seq data. RSEM (version 1.1.17) [91] with the default parameters was used to map the high-quality reads to the transcripts and calculate the expression levels (transcripts per million (TPM) and read count) of the protein-coding genes. Three replicates of each stage were used to reduce sampling bias. Differentially expressed genes among different developmental stages were detected with edgeR (version 3.30.3, the false discovery rate (FDR)<=0.05) [92] from the R package (version 3.6.1) using the read counts of the genes. Clustering of the gene expression time-course data from the four stages of *B. schroederi* was performed with Mfuzz (version 2.48.0) [93].

### Gene Ontology enrichment analysis

The tool BiNGO (Version 3.0.4) [94] implemented in Cytoscape (version 3.7.1) [95] software was used to analyze the GO enrichment of the genes from expanded gene families or differentially expressed genes with a hypergeometric test. The GO annotation profile of *B. schroederi* was constructed with an in-house Perl script, and the ontology file was obtained from the Gene Ontology web (http://geneontology.org/). GO terms with p-values<=0.05 calculated by hypergeometric test were extracted for functional analysis.

### Histological processing and analysis

The ascaridiod adults were fixed in formalin and routinely processed for histology as described elsewhere [7]. Briefly, fresh adults of *B. schroederi* (n=6), *A. suum* (n=8), *P. univalens* (n=8) and *T. canis* (n=8) as well as ursine *B. transfuga* (n=6) were fixed in 10% neutral phosphate-buffered formalin for 24 h. The portion (~1.5 cm) of the middle body of each fixed worm was then cut, oriented transversally and inserted into biopsy cassettes. All samples were washed in tap water and dehydrated by serial dilutions of alcohol (methyl, ethyl and absolute ethyl). Paraffin embedding was performed with a Leica Tissue Processing station using a 12-h protocol. Paraffin blocks were prepared using a Leica inclusion station. Afterward, each block was cut into 3-μm tissue sections using a Leica microtome. Each slide was stained with hematoxylin and eosin (HE). For each worm, three slides were examined under an inverted light microscope (Olympus FSX100, Olympus Corporation, Japan), and the cuticle thickness of these five ascaridiod species were then measured and compared. The data were expressed as the means ± standard deviations (SD). Comparisons between ascaridoid species were performed by one-way ANOVA, LSD and Scheffe’s test using SPSS (IBM SPSS Statistics for Windows, version 20; Armonk, NY: IBM Corp., USA). p values < 0.01 were considered to be significant.

### Identification of potential drug targets and vaccine candidates

All *B. schroederi* proteins were searched against lethal genes (WormBase WS226: WBPhenotype:0000050, WBPhenotype:0000054, WBPhenotype:0000060 and WBPhenotype:0000062 and subphenotypes), the ChEMBL database (https://www.ebi.ac.uk/chembl/) and host proteins (http://panda.genomics.org.cn/download.jsp) using BLASTP (E≤1×10^−10^). Genes with alignment length ratios and similarities higher than 0.5 were selected. The genes that are homologous to the lethal gene and ChEMBL database were then selected, the genes that are homologous to the host were removed, and single-copy genes were further screened. The following formula was used to assign a score to each potential drug target gene:

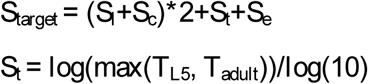

S_l_ and S_c_ are the SI values of genes homologous to the lethal gene and ChEMBL database, respectively, and T_L5_ and T_adult_ represent the gene expression (TPM) at the L5 and adult stages, respectively. S_e_ equals 1 if the target gene encodes a protease, protein kinase, protein phosphorylase, transporter or ion channel; otherwise, S_e_ is 0.

All proteins with signal peptides and one transmembrane structure domain were identified as E/S proteins. Proteases and protease inhibitors without host homology and TPM>=1 were screened from secretory proteins as vaccine candidates.

## Data Availability

All raw sequencing data (including the genome, transcriptome and Hi-C data) described in this manuscript have been deposited in the Sequence Read Archive (SRA) database under the accession codes PRJNA666314.

## Ethical statement

This study was approved by the Animal Ethics Committee of Sichuan Agricultural University (Sichuan, China; approval no. SYXK 2014-187) and the Wildlife Management and Animal Welfare Committee of China, and all procedures involving animals in the present study were in strict accordance with the Guide for the Care and Use of Laboratory Animals (National Research Council, Bethesda, MD, USA) and the recommendations in the ARRIVE guidelines (https://www.nc3rs.org.uk/arrive-guidelines).

## CRediT author statement

**Yue Xie:** Investigation, Formal analysis, Funding acquisition, Writing - original draft, Writing - review & editing. **Sen Wang:** Methodology, Formal analysis, Data Curation, Visualization, Software, Writing - original draft. **Shuangyang Wu:** Methodology, Formal analysis, Visualization, Writing - original draft. **Shenghan Gao:** Methodology, Formal analysis, Funding acquisition, Writing - review & editing. **Qingshu Meng**: Formal analysis, Writing - review & editing. **Chengdong Wang:** Resources. **Jingchao Lan:** Formal analysis. **Li Luo:** Resources. **Xuan Zhou:** Investigation. **Jing Xu:** Formal analysis. **Xiaobin Gu:** Resources. **Ran He:** Investigation. **Zijiang Yang:** Formal analysis. **Xuerong Peng:** Resources. **Songnian Hu:** Conceptualization, Supervision, Resources, Writing - review & editing. **Guangyou Yang:** Conceptualization, Supervision, Resources, Funding acquisition, Writing - review & editing. All authors read and approved the final manuscript.

## Competing interests

The authors have declared no competing interests.

## Acknowledgments

This research was partially supported by the Chengdu Giant Panda Breeding Research Foundation (Grant No. CPF-2012-13), Self-supporting Project of Chengdu Giant Panda Breeding Research Base (Grant No. 2020CPB-B20), Sichuan International Science and Technology Innovation Cooperation/Hong Kong, Macao and Taiwan Science and Technology Innovation Cooperation Project, China (Grant No. 2019YFH0065), Natural Science Foundation of China, Young Scientists Fund (Grant No. 31801048), and the High-level Scientific Research Foundation for the Introduction of Talents of Sichuan Agricultural University of China (Grant No. 03120322). The funders had no role in design, decision to publish, or preparation of the manuscript.

## Supplementary materials

**File S1 Supplementary note**

**Figure S1 Lifecycle of *B. schroederi* in the giant panda**

Fertilized eggs are excreted into the environment with feces. After one molt within the eggs, the embryos develop through first-stage larvae (L1s) to second-stage larvae (L2s). Host infections start with the oral intake of L2-containing infective eggs, and the L2s then hatch in the gastrointestinal tract and penetrate the intestinal walls of the host. The L2s are transported through the mesenterial blood veins to various organs (such as the liver and lungs) and induce visceral larva migrans (VLM). With further development to third-stage larvae (L3s), these larvae undergo hepatopulmonary and somatic migrations and are eventually swallowed again. In the small intestine, the third-stage larvae molt twice through the fourth and fifth stages and reach maturity. Dioecious adults mate, and the females can begin to excrete fertilized eggs again. It has been estimated that the lifecycle can be completed within three months. The adults can lead to abdominal pain, diarrhea, and potentially life-threatening intestinal blockage (IB) in giant pandas.

**Figure S2 Hi-C contact map of the *B. schroederi* genome with 27 chromosome-level pseudomolecules**

The signal represents two contact positions.

**Figure S3 Synteny comparison of *B. schroederi* and *A. suum***

The gray bands represent *A. suum* chromosomes, and the colorful bands note *B. schroederi* chromosomes. The links represent one-to-one orthologs between two species.

**Figure S4 Summary of tRNA gene content in *B. schroederi* and *C. elegans***

**A**. *B. schroederi* tRNA gene content and codon usage. **B**. The tRNA gene content and codon usage exhibit a high correlation in *B. schroederi*. **C**. *C. elegans* tRNA gene content and codon usage. **D**. The tRNA gene content and codon use show a high correlation in *C. elegans*.

**Figure S5 Venn diagram of homologous genes**

Venn diagram showing the number of orthologs between *B. schroederi* and three other ascaridoid species (*A. suum, P. univalens* and *T. canis*) after pairwise comparisons.

**Figure S6 Global phylogeny among nematode representatives of the phylum Nematoda with sequenced genomes**

The phylogenetic tree was constructed using concatenated amino acid sequences for 329 single-copy genes present in 12 genomes through maximum likelihood analysis. The numbers at the nodes indicate bootstrap values, and the number on each branch shows the distance.

**Figure S7 Estimation of the divergence time**

The times of divergence were estimated through an analysis of 1:1:1 orthologs between *B. schroederi* and 11 other nematodes, including three ascaridoid species, *A. suum, P. univalens* and *T. canis*. The distances are shown in million years ago (Mya).

**Figure S8 Ascaridoid orthologous gene similarity index**

**A**. Similarity index of orthologous genes of *T. canis* and *A. suum, P. univalens*, and *B. schroederi*. **B**.

Similarity index of orthologous genes of *B. schroederi* and *A. suum, P. univalens*. A higher similarity index indicates that the homologous genes are more similar. The p-value indicates whether the similarity difference between different species is significant.

**Figure S9 Cuticle thickness of ascaridoids**

**A**. Cuticle thickness of *B. schroederi* under a microscope (left: 40×, middle: 200×, right: 400×). Cuticle thickness of *A. suum* (**B**), *P. univalens* (**C**) and *T. canis* (**D**), respectively, under the microscope (left: 40×, middle: 200×, right: 400×).

**Figure S10 Cuticle thickness of *Baylisascaris transfuga***

Cuticle thickness of the congeneric *Baylisascaris transfuga* under the microscope at magnifications 40× (**A**), 100× (**B**) and 200× (**C**), respectively.

**Figure S11 Sequencing coverage of the region where collagens are tandem expansion genes**

The uniform coverage indicates that gene expansion exists and is not caused by assembly errors.

**Figure S12 Clustering analysis of gene expression in *B. schroederi* at four stages**

Twenty-five clusters were associated with the egg, L2, L5 and adult stages of *B. schroederi* based on the extent of shared genes among them, and each cluster was characterized primarily by a high level of gene expression at one of the four developmental stages.

**Figure S13 Genome survey of *B. schroederi***

The K-mer frequency was calculated with Jellyfish, and GenomeScope was used to estimate the genome size, heterozygosity and repeat content of *B. schroederi*.

**Supplementary Table 1 Statistics of the *B. schroederi* genomic sequencing data**

**Supplementary Table 2 Assessment of the quality of the *B. schroederi* draft genome**

**Supplementary Table 3 BUSCO analysis**

**Supplementary Table 4 Mapping rate of RNA-seq data against the *B. schroederi* genome**

**Supplementary Table 5 Summary of repeats in the *B. schroederi* genome**

**Supplementary Table 6 Statistics of tRNAs and rRNAs in the *B. schroederi* genome**

**Supplementary Table 7 Summary of tRNA genes in the *B. schroederi* genome**

**Supplementary Table 8 Statistics of gene models of *B. schroederi***

**Supplementary Table 9 Genes supported by RNA-seq data from *B. schroederi***

**Supplementary Table 10 Functional annotation of genes in *B. schroederi***

**Supplementary Table 11 Orthologous genes of *B. schroeder*i linked to one or more known KEGG pathways in *C. elegans***

**Supplementary Table 12 Antimicrobial effectors identified from *B. schroederi***

**Supplementary Table 13a Divergent genes among *B. schroederi, A. suum* and *P. univalens***

**Supplementary Table 13b GO enrichment of divergent genes**

**Supplementary Table 14 Genes inferred to be associated with host shift during the evolutionary history of ascaridoids**

**Supplementary Table 15a Expanded gene families of *B. schroederi***

**Supplementary Table 15b GO enrichment of genes from expanded gene families of *B. schroederi***

**Supplementary Table 16a Positively selected genes identified in *B. schroederi***

**Supplementary Table 16b GO enrichment of positively selected genes identified in *B. schroederi***

**Supplementary Table 17 Genes encoding nematode cuticle collagens and cuticlins identified in *B. schroederi***

**Supplementary Table 18 Comparison of the cuticles of ascaridoid species**

**Supplementary Table 19 GO enrichment of high expression genes in the different developmental stages of *B. schroederi***

**Supplementary Table 20 GO enrichment of down/up-regulated genes between developmental stages in *B. schroederi***

**Supplementary Table 21 Drug target candidates in *B. schroederi***

**Supplementary Table 22 Excretory/secretory (E/S) proteins of *B. schroederi***

**Supplementary Table 23 Vaccine candidates in *B. schroederi***

